# Serovar-level Identification of Bacterial Foodborne Pathogens From Full-length 16S rRNA Gene Sequencing

**DOI:** 10.1101/2023.06.28.546915

**Authors:** Dmitry Grinevich, Lyndy Harden, Siddhartha Thakur, Benjamin J Callahan

**Author notes:** Address correspondence to Benjamin Callahan.

## Abstract

The resolution of variation within species is critical for interpreting and acting on many microbial measurements. In the key foodborne pathogens *Escherichia coli* and *Salmonella*, the primary sub-species classification scheme used is serotyping: differentiating variants within these species by surface antigen profiles. Serotype prediction from whole-genome sequencing (WGS) of isolates is now seen as comparable or preferable to traditional laboratory methods where WGS is available. However, laboratory and WGS methods depend on an isolation step that is time-consuming and incompletely represents the sample when multiple strains are present. Community sequencing approaches that skip the isolation step are therefore of interest for pathogen surveillance. Here we evaluated the viability of amplicon sequencing of the full-length 16S rRNA gene for serotyping *S. enterica* and *E. coli*. We developed a novel algorithm for serotype prediction, implemented as an R package (Seroplacer), which takes as input full-length 16S rRNA gene sequences and outputs serovar predictions after phylogenetic placement into a reference phylogeny. We achieved over 89% accuracy in predicting *Salmonella* serotypes on *in silico* test data, and identified key pathogenic serovars of *Salmonella* and *E. coli* in isolate and environmental test samples. Although serotype prediction from 16S sequences is not as accurate as serotype prediction from WGS of isolates, the potential to identify dangerous serovars directly from amplicon sequencing of environmental samples is intriguing for pathogen surveillance. The capabilities developed here are also broadly relevant to other applications where intra-species variation and direct sequencing from environmental samples could be valuable.

## Introduction

Foodborne illnesses caused by bacterial pathogens are a major cause of hospitalizations and deaths throughout the United States, and the sub-species classification of different serovars within important enteric pathogens is important for surveillance and intervention. *Salmonella* species are the leading cause of hospitalizations according to CDC data, accounting for over 23,000 annual hospitalizations and over 450 annual deaths (Scallan et al. 2011). Treatment of salmonellosis has been estimated to cost over 2.5 billion dollars annually (Hoffman et al. 2012). The “big six” *E. coli* serotypes, the primary pathogenic *E. coli* causing foodborne illness, are responsible for nearly 170,000 cases of foodborne illnesses in the United States annually (Bertoldi et al. 2021). The burden caused by foodborne pathogens calls for modern surveillance tools that can rapidly identify these pathogens, prevent their introduction into the food system and people’s bodies, and rapidly trace outbreaks back to their source. However, many pathogen species can also be commensal organisms and occupy a normal environmental niche in the mammalian gut (Ramos et al. 2020, Kittana et al. 2018). Upwards of 90% of *E. coli* strains are predicted to be commensal; we need ways to distinguish which strains are dangerous for human health and which are not (Ramos et al. 2020). The most common subspecies classification of both *Salmonella* and *E. coli* is division into serovars based on their antigenic properties. The vast majority of illness caused by *Salmonella* derives from a small number of serovars including Enteritidis, Newport, Typhimurium, Monophasic Typhimurium, and Javiana. Similar patterns hold in pathogenic *E. coli*, with for example serovar O157:H7 being responsible for a large fraction of serious disease cases.

In clinical and applied microbiology, species-level identification has become standard, but increasingly we recognize the importance of sub-species classifications in order to effectively act on microbiological measurements. A wide variety of sub-species typing methods are employed across different contexts, including methods such as biological testing for antigenic gene features, molecular typing methods including PCR and microarray marker gene analyses, and computational typing approaches using whole genome sequencing data (Grimont et al. 2007, Wattiau et al. 2011, Zhang et al. 2019, Yoshida et al. 2016, Peterson et al. 2010). Serotyping through traditional microbiology methods is the most widely used method for sub-species classification of enteric pathogens, but is labor intensive and time consuming. Traditional serotyping requires lab access to many antisera which are necessary for testing for a variety of possible surface antigens. In the past decade, sequencing has played an increasingly important role in subspecies identification and it is now common to perform computational serotyping using whole genome sequencing (WGS) data from isolates of enteric pathogens. Popular tools for computational serotyping include the *Salmonella in Silico* Serotyping Resource (SISTR) and SeqSero2 (Zhang et al. 2019, Yoshida et al. 2016). In *E. coli,* tools such as SerotypeFinder and ECTyper have been developed for computational O and H antigen prediction (Joensen et al. 2015, Bessonov et al. 2021). Computational serotyping from WGS data achieves high accuracy, but important limitations remain including the cost of WGS and the need to isolate and culture individual microbes from potentially complex samples to obtain the necessary genetic material.

Here we develop an approach using sequences of the full-length 16S rRNA to classify *Salmonella* and *E. coli* to the serovar level. If serovar-level resolution could be obtained from 16S rRNA gene sequences alone, then it may be possible to improve pathogen surveillance by applying 16S rRNA gene amplicon sequencing directly to complex food-associated samples without requiring an isolation and culture step. Furthermore, the recent progress in long-read sequencing platforms has made amplicon sequencing of the full-length 16S rRNA gene increasingly affordable, accessible and accurate. Serovar prediction from a marker gene is fundamentally different from prediction from WGS data, as the 16S rRNA gene can only provide phylogenetic information, not direct information on the O and H antigen gene sequences. Thus we leverage a reference database of *Salmonella* and *E. coli* strains with known serotypes, fully enumerated complements of their multi-copy 16S rRNA genes, and an associated high-quality phylogeny linking the reference strains. We evaluated the viability of serotyping enteric bacteria using full-length 16S rRNA gene sequencing data, and developed user-friendly software for serovar prediction from such data in *Salmonella* and *E. coli*. Although our focus here is on *Salmonella* and *E. coli*, the methods we develop are generalizable to many other bacterial taxa for which sufficient numbers of sequenced genomes and appropriate sub-species classification schema are available.

## Methods

### Curation of Reference Database

Complete tables of all available reference assemblies were downloaded from the GenomeTrakr database (https://www.ncbi.nlm.nih.gov/pathogens/) for both *S. enterica* (July 10, 2022) and *E. coli* (September 5, 2022). Initial filtering was done by choosing all assemblies with assembly completion thresholds of “chromosome” or “complete genome”, as determined by NCBI. We also required that all reference assemblies contained exactly 7 full-length 16S rRNA gene sequences, and that whole genome computational serovar prediction yielded a complete and unambiguous serovar. For *Salmonella* entries, we removed reference assemblies which failed to receive an O-antigen assignment, identified as O-antigen assignments marked “-” in the output of SISTR. For *E. coli* entries, we removed reference assemblies which were missing either an O-or H-antigen assignment, identified as assignments marked as “-” in the output of ECTyper. The final reference database(s) were stored as a FASTA file, with 7 16S sequences for each reference genome labeled in the FASTA file by the original assembly, 16S rRNA allele number, and computational serovar assignment.

### Serovar Assignment to Reference Genomes

*In silico* serovar prediction on *Salmonella* assemblies was done using the *Salmonella In Silico* Typing Resources (SISTR ver. 1.1.1, available at https://github.com/phac-nml/sistr_cmd). SISTR was run using the sistr_cmd command following recommended standard parameters (sistr_cmd - o json -qc). Serovar prediction on *E. coli* was performed using the ECTyper tool (available at https://github.com/phac-nml/ecoli_serotyping). Analysis was run using the ectyper command with default options and a pre-sketched mash archive downloaded on August 19, 2021 from https://gembox.cbcb.umd.edu/mash/refseq.genomes.k21s1000.msh. The Typhimurium and Monophasic Typhimurium serovars in *Salmonella* form a single distinct clade that cannot be distinguished using core genes or 16S sequences. Thus, we chose to merge them in our reporting and classify them as a single group labeled “Typhimurium/Monophasic Typhimurium”.

### Construction of Reference Phylogenetic Trees

Core genes for each species were defined as those genes present in every reference genome. We identified a pool of 100 core genes for *Salmonella* and 101 core genes for *E. coli* species. MAFFT (ver. 7.490, available at https://mafft.cbrc.jp/alignment/software/) was used to generate multiple sequence alignments for each core gene (parameters: --retree 2 --inputorder). The concatenated alignments of all core genes were then used to infer the phylogeny linking the reference strains using the RAxML-ng tool (ver. 1.1.0, available at https://github.com/amkozlov/raxml-ng) with the raxml-ng command (parameters: –model GTR+G) (Stamatakis 2014, Kozlov et al. 2019). The final tree was stored in newick format for input into downstream phylogenetic placement analysis.

### Serovar Assignment

Our serovar assignment algorithm takes as input a set of 7 16S rRNA gene sequences corresponding to the 7 genomic copies of the 16S rRNA gene in both *Salmonella* and *E. coli*. Query sequences are first aligned to a pre-aligned 16S allele reference containing all of our reference 16S sequences using the mafft command from MAFFT (parameters: --add -- keeplength). The correct relative ordering of the query 16S sequences and the 16S sequences retrieved from the reference assembly is unknown *a priori*. So at this stage the set of reference 16S alleles are re-arranged to optimally match the query. More precisely, for each reference assembly in the database, its set of 7 16S alleles is compared against the 7 query sequences, and the reference alleles are rearranged into the order that minimizes total nucleotide mismatches to the query. Query sequences and re-arranged reference 16S sequences are concatenated and stored as FASTA input files for phylogenetic placement through EPA-NG (ver. 0.3.8, available at https://github.com/Pbdas/epa-ng) (Berger et al. 2011, Barbera et al. 2019). The query is phylogenetically placed into the reference phylogeny using epa-ng command (parameters: – filter-max 100) and a model parameter specification evaluated from RAXML-ng using the evaluate command (raxml-ng --evaluate --msa $REF_MSA --tree $TREE --prefix info --model GTR+G+F). After query sequence placement, all placements reported by EPA-NG within the 99.9% likelihood weight ratio (LWR) range are considered candidate placements.

Candidate placements of the query into the reference phylogeny are assigned to either the immediately distal or immediately proximal node to the insertion site of the pendant branch, whichever was closer to the insertion point. The initial MRCA is defined as the MRCA of the nodes assigned to each candidate placement. In order to account for uncertainty in the phylogenetic placements, the initial MRCA is then moved higher in the tree in a step we call “pendant length adjustment”. A branch length K is defined as the maximum pendant length from the candidate placements multiplied by an arbitrary scalar value. The phylogenetic tree is traversed from the initial MRCA towards the root by branch length K, and the closest node (distal or proximal) to this position is defined as the pendant-adjusted MRCA. If all candidate placements have short pendant lengths (suggestive of high confidence in the insertion site), it is possible that the MRCA will not change during pendant length adjustment. Here we used a pendant length multiplier value of 1.5 based on a data-driven evaluation in *Salmonella* and *E. coli*.

The set of reference assemblies contained in the clade defined by the pendant-adjusted MRCA is considered the set of “hits” of our algorithm. A specific serovar assignment is made if greater than some threshold fraction T of the hits are of the same serovar. By default, our computational tool sets T=50%, and reports serovar as indeterminate if no serovar exceeds 50% of the hits. However, users can also obtain a complete reporting of all serovars included in the final hits and their associated percentages.

### The *Seroplacer* R Package

The serovar prediction algorithm described above is available as an R package downloadable from Github (https://github.com/Dogrinev/Seroplacer). The input used by the algorithm is a FASTA file containing 7 16S ribosomal gene sequences. The *Data_Preparation* function re-formats the sequence data from the user input FASTA file for use in the placement algorithm. The serovar prediction method employed by the algorithm depends on two external command line tools, EPA-NG and MAFFT (Katoh et al. 2002, Berger et al. 2011, Barbera et al. 2019), which are executable through provided wrapper functions in the package, *mafft_wrap* and *epa_ng_wrap*. The 16S rRNA gene reference sequence data used for serovar classification are stored in non-aligned and pre-aligned FASTA files. The reference phylogenetic trees used by the package are stored in newick tree format. After phylogenetic placement through the core function *Clade_Hit_Finder_Pendant*, the software outputs a table containing all serovars identified in the final phylogenetic clade predicted, the associated percentages of each serovar, and a serovar prediction.

### Environmental and Sample Isolates

We obtained 10 *Salmonella enterica* isolates from previously collected stocks obtained during pathogen surveillance by the Thakur Molecular Epidemiology Laboratory at North Carolina State University. These isolates covered a variety of isolation sources including human feces, vegetative buffer, sheep feces, lairage swabs, caprine feces, and cecal content. We obtained 27 animal samples from routine retail meat surveillance testing that contained a variety of microbes with a fraction of samples containing *Escherichia coli*. Meat samples were enriched in buffered peptone water and stored for sequencing analysis.

### Environmental Isolate Sequencing

Genomic DNA for the bacterial isolates was extracted using the DNeasy PowerLyser Microbial Kit following manufacturer protocols (Qiagen). DNA concentrations were measured using NanoDrop Spectrophotometry (Thermo Fisher Scientific) and normalized to contain approximately 100 ng/µL and 30 µL of bacterial DNA. 16S sequencing libraries for 10 *Salmonella* isolates and 27 animal sourced isolates were prepared using the LoopSeq pipeline from Loop Genomics following manufacturer protocols (loopgenomics.com). Loop Genomics 16S primers were used to amplify the specific 16S region (Forward: AGAGTTTGATCMTGGC, Reverse: TACCTTGTTACGACTT). Isolate samples were prepared to measure 1000 ∼1.5kb molecules per sample, and environmental samples were prepared to measure 50000 ∼1.5kb molecules per sample. Amplicon sequencing data was quality filtered, denoised, and analyzed using primarily the standard dada2 workflow (available at https://github.com/benjjneb/dada2) with modified parameters designed to improve sensitivity when analyzing LoopSeq data (Callahan et al. 2021). Adjustments included changes to the filterAndTrim function (maxEE=0.5) and the dada function (OMEGA_A=1e-10, DETECT_SINGLETONS=TRUE). Denoised 16S sequences were assembled into sets of 7 16S gene alleles and converted to FASTA file format for serovar analysis. 16S query sequences were inputted into the serovar placement algorithm described previously and final serovar predictions were recorded.

### Computational Analyses and Statistics

Statistical analyses and visualizations were performed using R statistical computing software (version 4.2.1) and RStudio (version 2022.07.1+554). Phylogenetic distance calculations throughout this work are all performed using the “ape” package (ver. 5.6-2) in R.

### Data Availability

Reproducible analysis of computational code and generation of figures and data can be viewed at https://github.com/Dogrinev/Seroplacer_Manuscript. Analyses are available in the form of an HTML containing all outputs, or R markdown files to re-run all code. Raw sequence data for environmental samples and bacterial isolates analyzed by amplicon sequencing can be found in the Sequence Read Archive (SRA) available at Bioproject ID PRJNA988246.

## Results

### Curation of Reference Sequence Data and Phylogenetic Trees

We downloaded a list of 1,787 *Salmonella* assemblies with “chromosome” or “complete genome” quality from the GenomeTrakr database (https://www.ncbi.nlm.nih.gov/pathogens/) on July 10, 2022. Of these, 1,700 reference assemblies contained exactly 7 full-length 16S rRNA gene sequences and a complete set of *Salmonella* core genes. We used the SISTR serotyping method to assign serovars to each reference assembly (Yoshida et al. 2016). 82 entries were not assigned an O-antigen label, and were removed, leaving 1,618 assemblies in our final *Salmonella* reference database representing 155 unique serotypes. A phylogenetic tree was inferred from a concatenated alignment of the *Salmonella* core genes using RaXML and 20 bootstrapped replicates (Methods). Our phylogeny recapitulated known major features of the *Salmonella* phylogeny, including reconstructing known clade groups (I-V). All 7 full-length 16S rRNA gene sequences were extracted from each reference assembly, and organized into two reference FASTA files, one in which the sequences are unaligned and another in which they are aligned using MAFFT (Methods). The ID line of the reference fasta files were formatted with the assembly identifier, the 16S gene copy number (1-7), and the assigned serotype. We followed the same process to develop a reference database for *E. coli*, yielding a total of 2,791 assemblies representing 692 unique serotypes in our *E. coli* reference database.

## 16S rRNA Gene Profiles are On Average More Similar Within Serovars

We performed an initial evaluation of whether 16S rRNA gene sequences carry sufficient signal to differentiate between serovars by comparing the average dissimilarity in 16S rRNA gene profiles between and within *Salmonella* serovars. For clarity, the “16S rRNA gene profile” refers to the full complement of 16S rRNA gene alleles from a given genome, in arbitrary order. Here that means 16S rRNA gene profiles consist of 7 sequences, as both *Salmonella* and *E. coli* have 7 copies of the 16S rRNA gene in their genomes. We calculated the number of differing nucleotide sites between alignments of the 16S rRNA gene allele profile present in each *Salmonella* genome (Methods), and then averaged these pairwise nucleotide dissimilarities between and within each serovar with at least 4 entries in our reference database (Figure 2).

**Figure 1.**
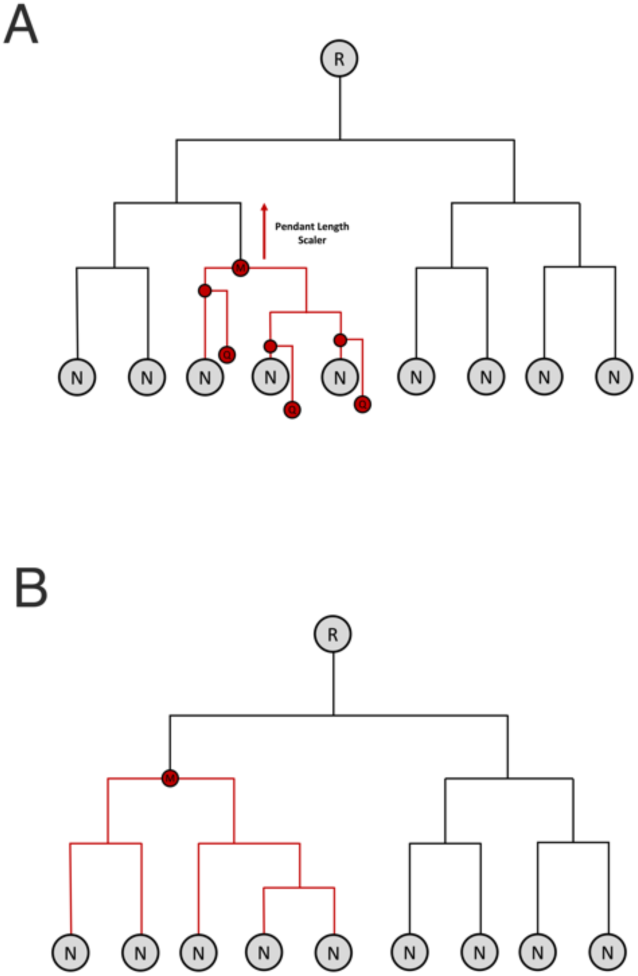
Proposed serovar assignment algorithm. A) Queries (Q) consisting of the full complement of 7 16S rRNA gene sequences are phylogenetically placed into a reference phylogenetic tree consisting of terminal nodes (N) and root (R). Typically, multiple potential placements are predicted, from which an initial most-recent-common ancestor (MRCA; M) is constructed. The maximum pendant length (i.e. the length of the new branch inserted into the tree to place the query) is recorded. The MRCA is then adjusted upwards towards the root of the tree by the maximum pendant length times a predetermined scalar value of order 1, yielding the final MRCA shown in (B). A serovar assignment is made if more than a threshold fraction (by default 0.5) of the reference entries contained in the phylogenetic clade defined by the final MRCA are of the same serovar.

**Figure 2.**
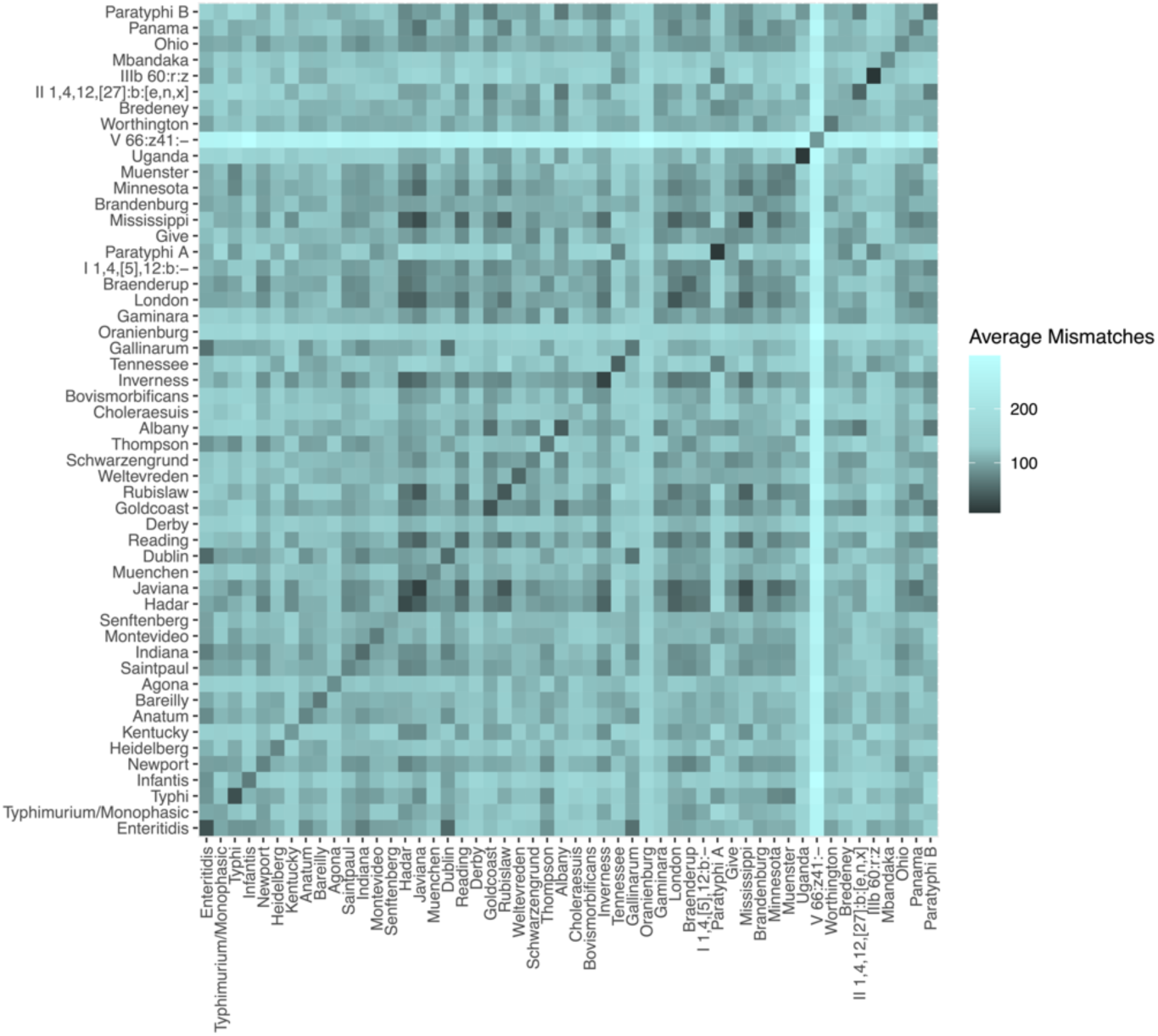
Average dissimilarity between the 16S rRNA gene profiles of *Salmonella* serovars. Pairwise dissimilarities were defined as the number of nucleotide mismatches between alignments of the full set of 16S rRNA genes (the 16S rRNA gene profile) of *Salmonella* assemblies, after optimally re-arranging the profiles relative to one another (Methods). The dissimilarity between assemblies of the same serovar was lower on average than between different serovars (darker colors on diagonal). However, some pairs of serovars had 16S rRNA gene profiles that were as or more similar than were the profiles within those serovars (dark cells off the diagonal).

*Salmonella* strains of the same serovar have, on average, lower dissimilarity across their 16S rRNA gene profile than do *Salmonella* strains of different serovars (darker colors along diagonal, Figure 2), but not always. The average number of nucleotide mismatches is typically greater than 100 (>0.1% dissimilarity) when comparing the 16S rRNA gene profiles from different *Salmonella* serovars. Profiles from same-serovar strains show lower numbers of nucleotide mismatches, with typical values in the 30-50 range (<0.05% dissimilarity). The higher similarity of 16S profiles from same-serovar strains suggests that 16S sequences may carry enough signal to make serovar assignments in some cases. However, we also observed low values (<0.05% dissimilarity) between some different pairs of serovars (dark colors off the diagonal, Figure 2). Thus, 16S rRNA gene profile similarity alone may not always provide sufficient resolution to discriminate serovars. *E. coli* had similar overall patterns: Same-serovar *E. coli* strains showed average nucleotide mismatch values in the 40-70 range, with higher dissimilarity usually, but not always, when comparing across serovars (Supplemental Figure 1).

### *Salmonella* Serovars are Phylogenetically Coherent, but Sometimes Polyphyletic

All large *Salmonella* serovars (20+ assemblies in our reference database) were coherently phylogenetically organized, but sometimes into more than one clade. The three largest serovars in our phylogenetic tree, including Enteritidis, Typhi, and Typhimurium, clustered into single clades in which the vast majority of assemblies were assigned to the same serovar (Figure 3A, Typhi). However, other serovars clustered into two or even three distinct clades, as can be observed in *Salmonella* serovar Kentucky (Figure 3B). The polyphyletic structure of some serovars complicates the use of simple similarity-based methods for serovar identification, but is not in principle problematic for methods based on phylogenetic placement. Out of 21 *E. coli* serovars with the largest representation in our reference database, 14 clustered into single coherent clades, including the three largest serovars (O157:H7, O16:H48, and O25:H4). Polyphyletic organization into two distinct clades was observed in 6 large serovars, while 1 large serovar (O11:H25) only partially clustered with a fraction of its assemblies spread throughout the phylogenetic tree.

**Figure 3.**
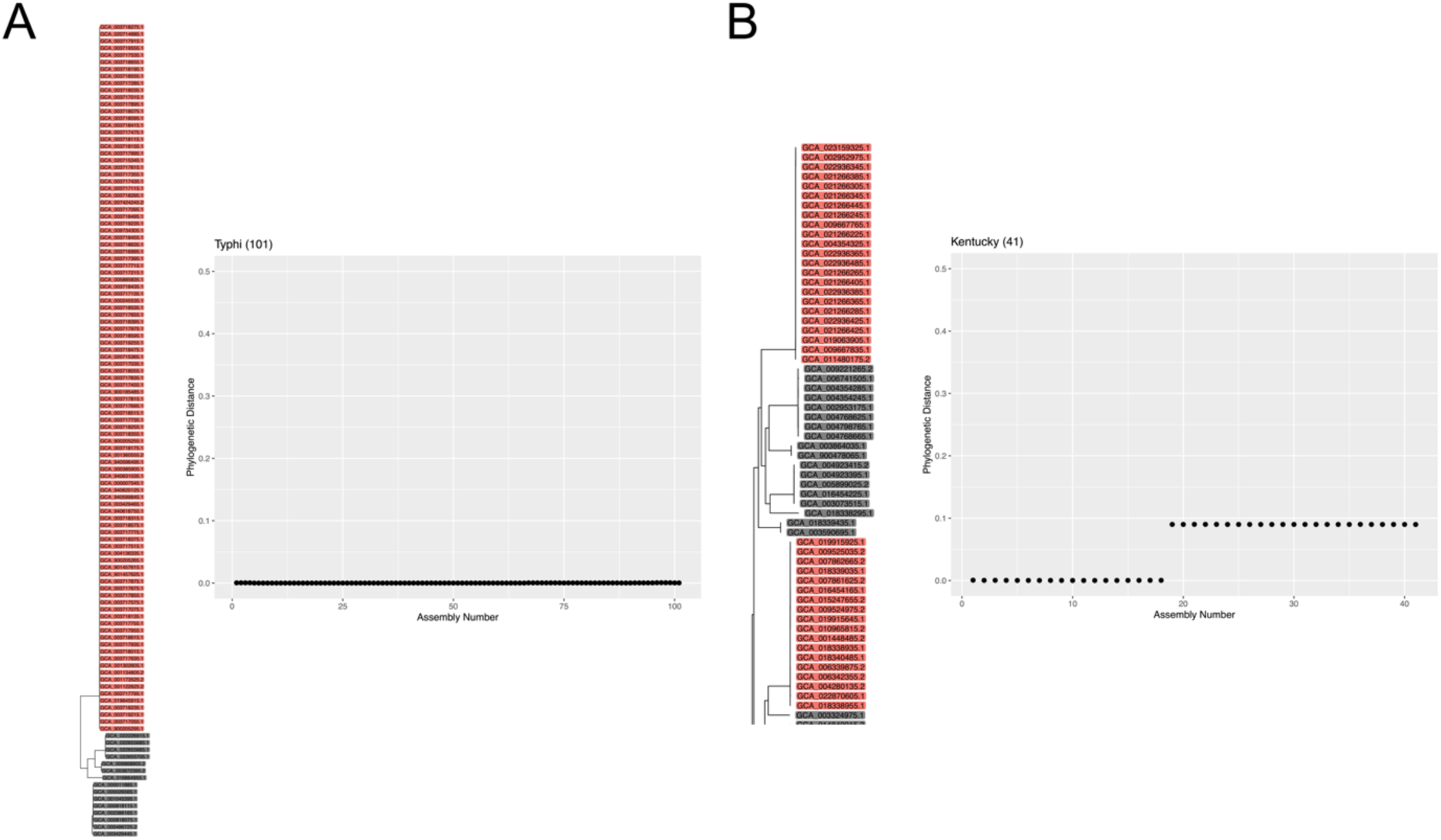
Monophyletic and polyphyletic *Salmonella* serovars. **(A)** Phylogenetic distances between an example Typhi assembly and all other Typhi assemblies in our *Salmonella* reference database. The portion of the phylogenetic tree containing all Typhi assemblies shows they are closely related and form a coherent clade. **(B)** Phylogenetic distances between an example Kentucky assembly and all other Kentucky assemblies in our *Salmonella* reference database. Kentucky strains form two distinct clades within the phylogenetic tree. Some assemblies in the reference phylogeny were removed in panel B for visual clarity.

### Method and Software for Serovar Assignment From 16S rRNA Gene Profiles

We developed a new algorithm for predicting serovar from 16S rRNA gene profiles based on phylogenetic placement into a reference phylogeny of serovar-labeled reference assemblies (see Methods for details). In short, we first curated reference databases of high-quality complete genomes with unambiguous serovar assignments for both *Salmonella* and *E. coli* (15). A core genome phylogeny for each reference database was constructed, and a reference fasta file with entries for each 16S rRNA gene allele from each reference assembly created. Queries, in the form of complete 16S rRNA gene profiles were optimally re-ordered and aligned to the reference 16S rRNA gene profiles. Concatenated alignments were used as input to the maximum-likelihood evolutionary placement algorithm (EPA-NG) to place query profiles into the reference phylogenetic tree. Phylogenetic placements and their associated uncertainty were used to define a set of “hits” in the reference database, and to make a serovar prediction. The full pipeline is organized as an R software package available at https://github.com/Dogrinev/Seroplacer.

### *In silico* Evaluation of Serovar Assignment from Full-length 16S rRNA Gene Sequencing

We performed an *in silico* evaluation of serovar assignment accuracy using a “leave-one-out” strategy, in which each entry in the reference database was first removed from the database, and then used as a query to our algorithm. That is, for each reference assembly, we deleted it from our reference database, and then used our method to classify the 16S rRNA gene profile of the deleted reference. We categorized our serovar prediction results into three groups: correct, incorrect, and indeterminate (see Methods for more details). Correct: A definite serovar assignment was made by our method that matched the ground truth serovar derived from whole-genome-sequencing (WGS). Indeterminate: No serovar assignment was made by our method because no single serovar represented a consensus (i.e. 50% or more) of all leaves within the final clade. Incorrect: A definite serovar assignment was made by our method that was different than the WGS serovar prediction. We developed two additional evaluation metrics to quantify the performance of our method: serovar accuracy and query recovery. Serovar accuracy (SA) is the fraction of hits under the final clade which correctly matched the serovar of the query. Query recovery (QR) is a binary value that is defined as TRUE when the final clade contains the attachment point of the query assembles phylogenetic branch (i.e. would contain the query if the query had not been removed from the reference), and FALSE otherwise.

Our algorithm incorporates uncertainty in phylogenetic placement to broaden the set of references considered as potential “hits”, and thus as contributors to the prediction of serovar. In a phylogenetic placement algorithm like EPA-NG, the pendant length represents the length of the newly inserted branch during phylogenetic placement into the existing tree. We considered this as a proxy for the uncertainty in the phylogenetic placement. More precisely, we adjusted the final prediction clade by traversing the reference tree upwards by a multiple of the maximum pendant length of qualified phylogenetic placements of the query (Methods). We tested a range of multipliers for the pendant length adjustment to identify the best value for this algorithm parameter. A larger pendant length adjustment makes the prediction clade more inclusive by shifting the MRCA upwards towards the root. An ideal pendant length multiplier value will produce prediction clades that contain the query strain (high QR), while not sacrificing too much specificity in the final result (high SA).

### Algorithm Performance in *Salmonella*

In *Salmonella*, our method had mean serovar accuracy between 83 and 91% across the range of pendant-length multiplier parameters considered (0.001x-8x) on our test dataset consisting of 52 Salmonella serovars with at least four entries in our reference database (Figure 4). As expected, positive predictive accuracy decreased as the pendant length multiplier was raised, because as the final clade grows in size, an increased number of queries returned indeterminate serovar assignments. However, this decrease in accuracy was minor for multipliers up to ∼4x. The fraction of queries recovered increased as the pendant length multiplier was raised, since larger final clades were more likely to contain the phylogenetic attachment point of the removed query. From these results, we selected a pendant length multiplier of 1.5 as providing a good balance of mean serovar accuracy (>89%) and mean query recovery (>88%, Figure 4).

**Figure 4.**
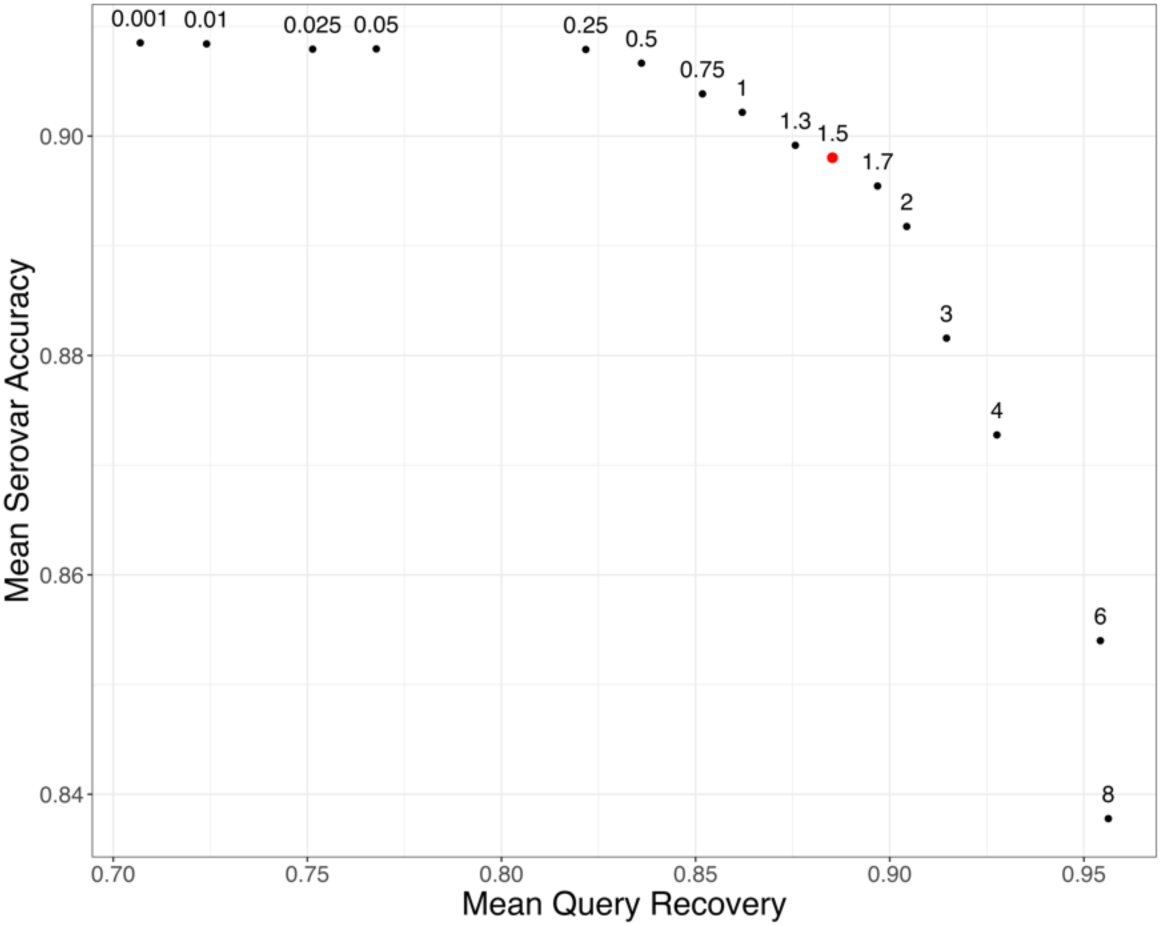
Serovar Placement Algorithm Performance on *Salmonella* Test Data. For each test query, a final clade of hits was determined by identifying the most recent common ancestor (MRCA) of all placements. The summary statistics plotted here are averages across all test queries for each given pendant length multiplier. Mean serovar accuracy represents the fraction of hits which match to the original query serovar averaged across all test queries. Mean query recovery is the fraction of original test queries which were located within the final calculated clade of hits. Pendant length multiplier values are labeled above the respective data point.

The accuracy of our serovar assignment method varied among *Salmonella* serovars. The fraction of different serovar assignment outcomes across the 52 serovars present in our test dataset was 90.1% correct; 4.5% incorrect, and 5.4% indeterminate, but clear differences among specific serovars were apparent (Figure 5). Some variation in serovar prediction accuracy is explained by the clear trend of decreased classification accuracy as the number of reference assemblies for a serovar decreased (Figure 5, Table 1). The largest serovars (Enteritidis: n=300, Typhimurium/Monophasic Typhimurium: n=288, Typhi: n=101) were all highly predictable (>90% accuracy). The proportion of correct assignments only began to drop below 75% in serovars containing fewer than 10 entries in the reference database. We calculated per-serovar *in silico* assignment outcomes for all serovars with at least four entries in our reference database (Supplemental Table 1).

**Figure 5.**
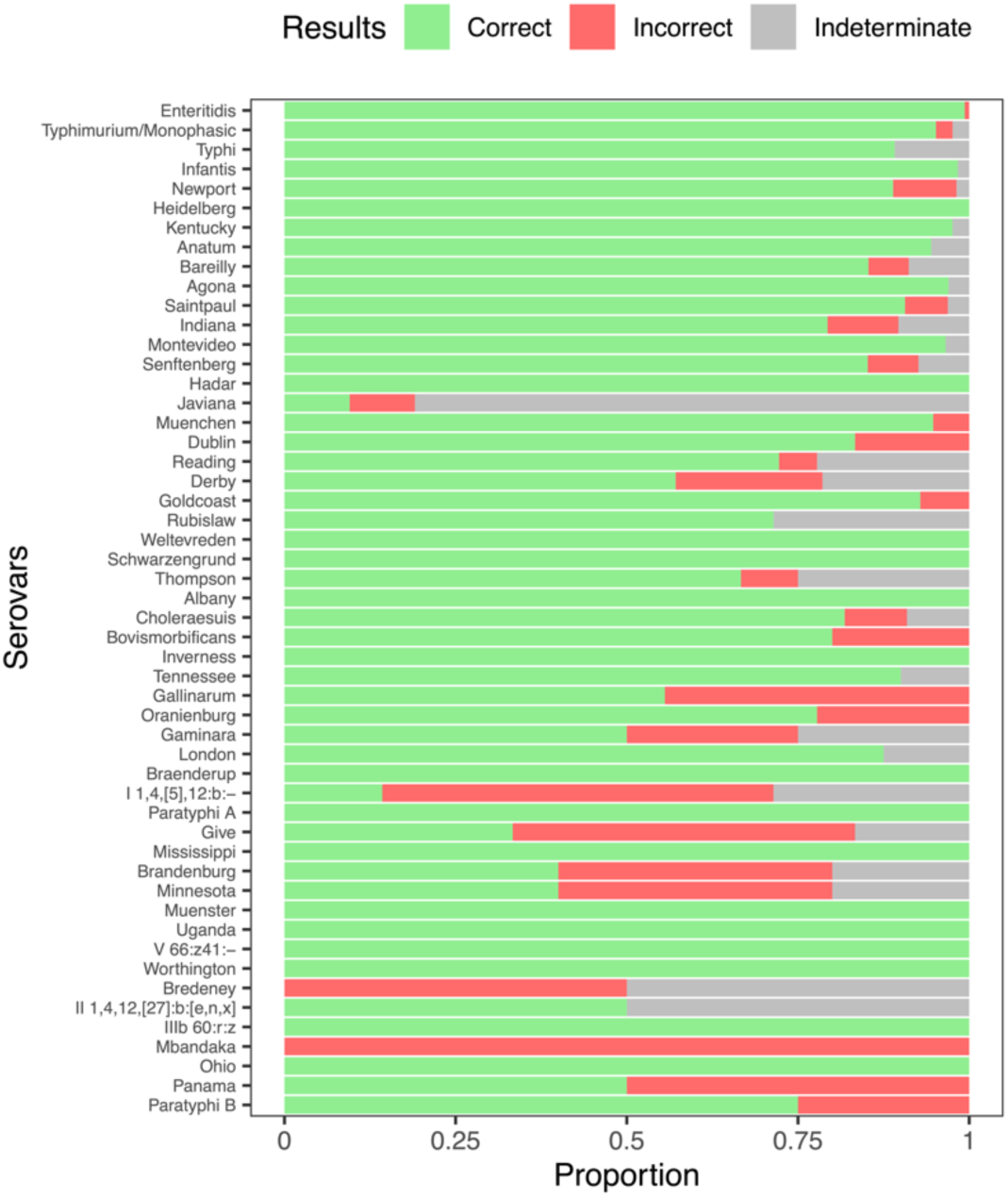
Serovar prediction accuracy in *Salmonella* varies across serovars. The proportion of query sequences that resulted in Correct, Incorrect or Indeterminate (Methods) serovar assignments from the 52 serovars with at least four entries in our *Salmonella* reference database. Serovars are ordered from most representation (Enteritidis, 300 assemblies) to least representation (Paratyphi B; 4 assemblies).

**Figure 6.**
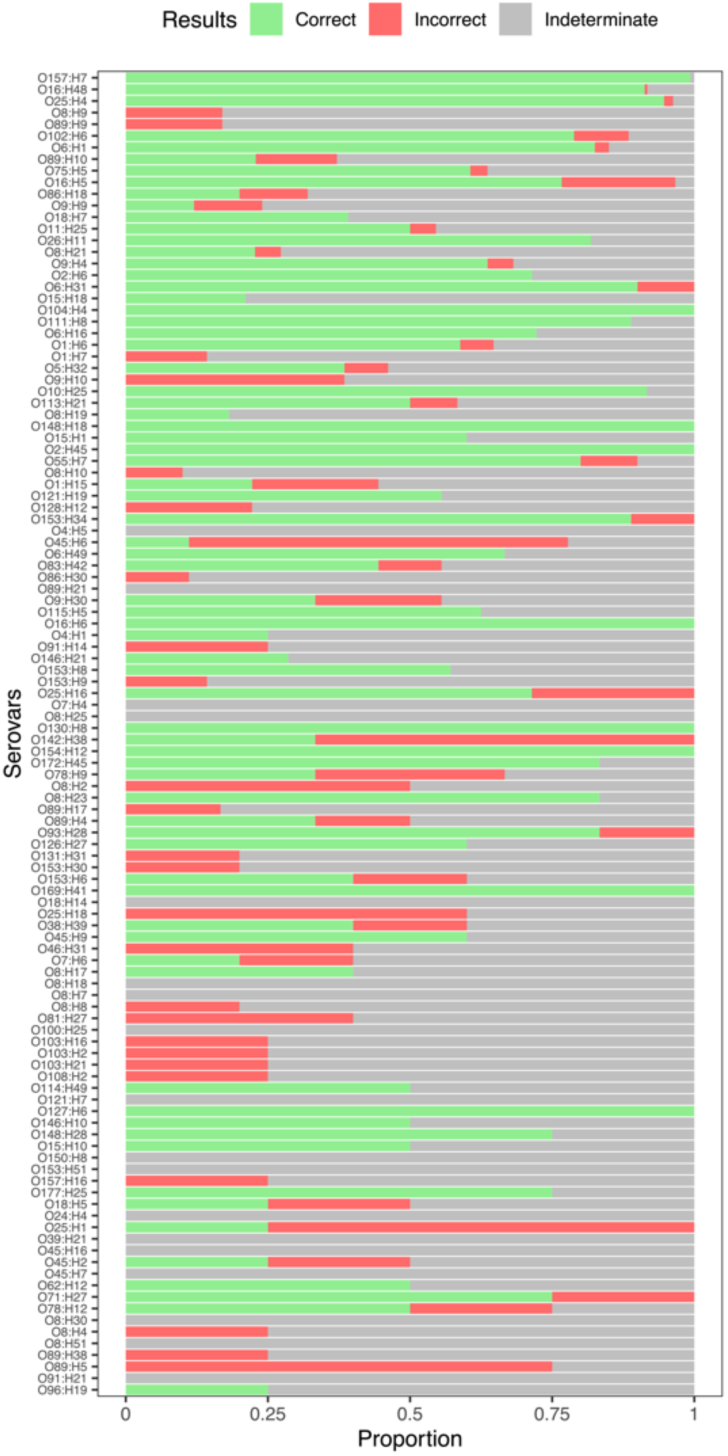
Serovar prediction accuracy in E. coli varies across serovars. The proportion of query sequences that resulted in Correct, Incorrect or Indeterminate (Methods) serovar assignments from the 114 serovars with at least four entries in our *E. coli* reference database. Serovars are ordered from most representation (O157:H7, 284 assemblies) to least representation (O96:H19; 4 assemblies). Assemblies with ambiguously assigned O antigens were excluded from this figure. Rates of indeterminate serovar assignment were much higher in *E. coli* than in *Salmonella*.

**Table 1.** Serovar assignment outcomes in *Salmonella* serovars depend on their level of representation in our reference database. Percentage correct, incorrect, and indeterminate rates are averages across all serovars in each reference-size category.

|  | *Correct* | *Incorrect* | *Indeterminate* |
| --- | --- | --- | --- |
| 100 - 300 Assemblies | 96.1% | 2.6% | 1.3% |
| 20 - 99 Assemblies | 89.4% | 7.1% | 3.5% |
| 10 - 19 Assemblies | 84.5% | 8.6% | 6.9% |
| 4 - 9 Assemblies | 68.0% | 9.6% | 22.4% |

Our method’s performance did not meaningfully degrade when identifying polyphyletic serovars. Some serovars such as Kentucky form two or more distinct clades within the *Salmonella* phylogeny (Figure 3). In the 20 *Salmonella* serovars with the largest representation in our reference database, we determined that 9 did not cluster into one clade. For 6 of these 9 large polyphyletic serovars (Newport, Kentucky, Bareilly, Saintpaul, Montevideo, and Muenchen) we still achieved over an 80% correct prediction rate, suggesting that it is not necessary for each reference to cluster perfectly into one clade in order to make good predictions. For the remaining 3 large polyphyletic serovars, we observed lower correct prediction accuracy (Reading 72%, Derby 57%, and Javiana 9%). Overall, these accuracy results are similar to those observed across comparably well-represented monophyletic serovars.

### Algorithm Performance in *E. coli*

We similarly evaluated our serovar prediction method in *E. coli*, and found that it was effective for some serovars but had a much higher rate of indeterminate assignments and a larger dropoff in accuracy for less-represented serovars than we observed in *Salmonella*. In the *E. coli* serovars with the most representation in our reference database (100+ assemblies), we made correct serovar calls in 96% of test queries, with a 2% indeterminate rate. After that, the rate of correct predictions dropped substantially. In moderately represented serovars (20-100 assemblies in the reference) our correct prediction rate was 47% with a 44.6% indeterminate rate. In serovars with 10-19 assemblies in the reference our correct prediction rate was 45.8% with a 46.5% indeterminate rate. Serovar prediction accuracy was worst in the serovars with the least representation in the reference database (4-9 assemblies). In this group, the method had a correct prediction accuracy of 29.7% with a 57.4% indeterminate rate. Incorrect predictions remained low to modest (<14%) across all reference-size groups (Table 2).

**Table 2.**
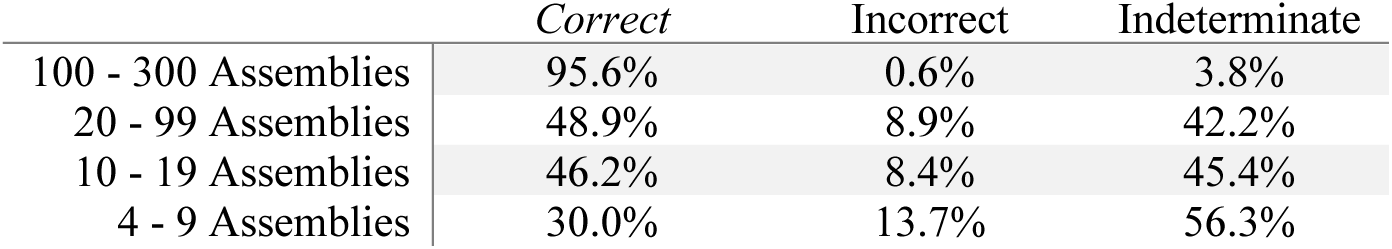
Serovar prediction accuracy in *E. coli* serovars depends on the amount of representation in our reference database. Percentage correct, incorrect, and indeterminate rates are averages across all serovars in a reference-size category.

**Table 3.** Results of serovar prediction on *Salmonella* bacterial isolates. We performed full-length 16S rRNA gene amplicon sequencing on a set of isolates for which whole genome sequencing (WGS) was already available. We report the prediction from our algorithm, the WGS serovar prediction, and the result of our method’s prediction treating the WGS prediction as the ground truth.

| Our Prediction<br>(% in Clade) | WGS Prediction | Result | Source | Accession |
| --- | --- | --- | --- | --- |
| Enteritidis (99%) | Enteritidis | Correct | Human Stool | SAMN11478315 |
| Enteritidis (99%) | Enteritidis | Correct | Not Listed | SAMN15508318 |
| Typhimurium/Monophasic<br>(100%) | Typhimurium | Correct | Vegetative<br>Buffer | PDT000640020.1 |
| Typhimurium/Monophasic<br>(99%) | Typhimurium | Correct | Human Stool | PDT000465258.1 |
| Enteritidis (99%) | Agona | Incorrect | Human Stool | PDT000550769.1 |
| Indeterminate | Muenster | Indeterminate | Sheep Feces | PDT000688730.1 |
| Typhimurium/Monophasic<br>(99%) | Monophasic<br>Typhimurium | Correct | Lairage Swab | SAMN13663197 |
| Indeterminate | Sundsvall | Not In Tree | Caprine Feces | SAMN13513401 |
| Indeterminate | Typhimurium | Indeterminate | Caprine Cecal<br>Content | PDT000641676.1 |
| Quebec/Soahanina<br>(50%/50%) | Sundsvall | Not In Tree | Caprine Feces | PDT000641787.1 |

In order to understand the difference in observed accuracy between *Salmonella* and *E. coli* serovar predictions, we further investigated the diversity and phylogenetic structure of serovars in each species. The *Salmonella* phylogeny contained 52 serovars containing at least 4 assemblies, while the *E. coli* phylogenetic tree contained 124 serovars containing at least 4 assemblies. It is possible that the increased variety of serovars in *E. coli* in combination with the much larger total phylogenetic tree makes it more difficult to make accurate serovar predictions. However, we suspect that *E. coli* serovars that do not form phylogenetically coherent clade(s) may also contribute to lower overall performance. In large *E. coli* serovars (20+ reference assemblies in our phylogenetic tree), we observed a polyphyletic clade structure in 6 serovars. Among these six large polyphyletic *E. coli* serovars, we had a poor prediction rate (<30% correct) in four: O8:H9, O89:H9, O9:H9, and O8:H21.

The serovar prediction accuracy of our method varied substantially among *E. coli* serovars, with notable differences in accuracy among O157 and the other “big six” *E. coli* serovars especially relevant to foodborne disease. Our method had a very high correct prediction rate (99.3%) in the highly relevant *E. coli* pathogenic serovar O157:H7. Our method also had a high correct prediction rate in O26 (81.8%) and O111 (88.9%). The correct prediction rate was lower for O45:H9 (60%) and O121:H19 (55.6%), and our method did not effectively predict (0%) in O103. There were no O145 reference entries in our dataset.

### Serovar Prediction on Environmental Samples

We evaluated the performance of our method on a set of *Salmonella* bacterial isolates collected from various environmental sources including human stool, animal waste, and lairage swabs. We performed amplicon sequencing of the full-length 16S rRNA gene and whole genome sequencing on 10 isolates obtained from environmental samples. In each sample, we identified a full profile of 7 16S alleles that was then used by our method to predict the serovar. We also made serovar predictions from the whole-genomes sequences as a ground truth (Methods). For 5 of the 10 environmental isolates, identical serovars were predicted based on our method and WGS. In three other isolates from serovars that were present in our reference database, our 16S-based method made two indeterminate predictions and one incorrect prediction. Two isolates were not able to be predicted with our method because the WGS predicted serovar (Sundsvall) was not present in our reference data. Notably, we made accurate predictions for the highly relevant foodborne pathogen serovars enteritidis, typhimurium, and monophasic typhimurium.

We also performed amplicon sequencing of the full-length 16S rRNA gene directly on 24 samples collected from retail meat rinsates, i.e. without first obtaining isolates. Of these 24 samples, 8 samples contained significant abundances of amplicon sequencing variants (ASVs) assigned to *E. coli* species, and 2 samples contained traces of *E. coli* but did not have sufficient sequencing depth to identify more than 1 ASV. *Salmonella* was not detected in any of the samples. We were able to clearly determine the full 7-allele rRNA gene profile in 3 of the 8 samples with significant *E. coli* abundance. We ran serovar predictions using our algorithm on these 3 samples and predicted that one sample contained *E. coli* serovar 0157:H7 and the other 2 resulted in an indeterminate result. The remaining 5 samples with significant *E. coli* abundances appeared to contain multiple strains in comparable abundances, and we were unable to unambiguously deconvolve the different strains’ 16S rRNA gene profiles.

## Discussion

Here we presented a computational tool that can make serovar-level assignments in *Salmonella* and *E. coli* from full-length 16S rRNA gene sequence data. Conventional wisdom might state that 16S rRNA gene sequencing can only achieve species-level resolution, but we think our results here make clear that sub-species resolution is possible from the kind of full-length high-accuracy sequencing now available. Unique aspects of our approach that enabled taxonomic resolution well below species-level were our utilization of the full set of alleles from the multi-copy 16S rRNA gene, and the use of phylogenetic placement into a serovar-labeled reference database. Our method was accurate for prediction of *Salmonella* serovars, was less accurate in *E. coli* generally but showed potential for prediction of pathogenic *E. coli* serovars from the “big six”, and is in principle generalizable to any other bacterial species. Due to the phylogenetic nature of our prediction algorithm, it depends on high-quality and extensive genomic reference sequence data to construct appropriate references for the bacterial species of interest. *Salmonella* and *E. coli* were a good starting point to develop our method because of the large quantity of reference data, with over 500,000 reference entries in *Salmonella* and 300,000 entries in *E. coli* available for download from the GenomeTrakr database. Our approach can easily be translated to other bacterial species with phylogenetically coherent sub-species classification schema by constructing a new phylogenetic tree for the organism of interest and assembling a reference 16S sequence database. Potential candidate bacteria for future application include *Staphylococcus aureus, Campylobacter jejuni, Klebsiella pneumoniae,* and *Listeria monocytogenes*.

The number of high-quality reference genomes available for a serovar was strongly correlated with our ability to make accurate predictions. Our method achieved its highest accuracies in serovars with large representation in our database for both *Salmonella* and *E. coli* (100+ assemblies). It is important to note that the number of genomes that could serve as effective references for our method was much smaller than the raw number present in many WGS databases, because we required genomes that fully resolved the multiple copies of the ribosomal operon. Resolving repeat elements like the rrn operon is a classic challenge for short-read sequencing, and most of the assemblies that qualified for our reference databases included long-read sequencing data. The GenomeTrakr database is being rapidly expanded, with over 5,000 new entries being submitted every month, but the rate of increase in high-quality genomes with resolved repeat elements is much smaller.

To our knowledge, our method is the first one that leverages the full complement of alleles from the 16S rRNA marker gene to improve the resolution of taxonomic classification, although some previous studies have provided case-study demonstrations (Callahan 2019; Ibal 2019; Johnson 2019). Amplicon sequencing randomly samples from the multiple copies of the 16S rRNA gene present in bacterial genomes, and variation between those alleles is increasingly evident when sequencing the full-length gene. Instead of considering this allelic variation just a nuisance, we chose to take advantage of this information by leveraging the data from the full set of 16S sequences for classification. It was common for two serovars to share a portion of identical 16S alleles but have unique differences in other alleles. Our method considers the unique nucleotide differences on a per-allele basis, meaning we can distinguish very similar isolates with nucleotide changes in specific 16S copies. The accuracy and length of amplicon sequencing with modern long-read sequencing platforms like Pacbio HiFi and Element Biosciences LoopSeq make this approach feasible (Callahan 2019; Karst 2021; Callahan 2021).

Evaluating whether a taxonomic classification method is of sufficient accuracy for specific applications is an important but difficult question that probably needs to meaningfully consider accuracy metrics on a per-taxon basis (Kralj 2020). Our method was only modestly accurate for predicting *E. coli* serovars in general: Roughly 40% of *in silico* test queries returned an indeterminate serovar classification, and 2/3 of the community samples for which we extracted an *E. coli* 16S rRNA gene profile also resulted in indeterminate classifications. However, the classification accuracy for the notorious O157:H7 serovar was exceptionally high for *in silico* testing, and O157:H7 was positively identified directly from amplicon sequencing in one of the community samples. If a specific application can accept a high rate of indeterminate serovar assignments for other *E. coli* serovars, but needs sensitive and specific detection of O157:H7, our method may be appropriate. More generally, if there are a defined set of species or sub-species taxa whose identification is critical to the mission, then accuracy of classification methods should focus primarily on those taxa over broad cross-taxa averages (Kralj 2020).

A weakness of our method and its envisaged application to amplicon sequencing from complex samples is its dependence on having the complete 16S rRNA gene profile as a query. This weakness is most acute in complex environmental samples, where strain confusion and low relative abundance can hamper the recovery of the full 16S rRNA gene profile from specific strains present in the sample. For example, in only 3/8 of the environmental samples considered here with significant *E. coli* relative abundance could we unambiguously determine a single strain profile from the mixture of *E. coli* 16S rRNA gene sequences. Advancing our approach to work with partial queries containing only some of a strain’s alleles is one path for improvement. Technological improvements in the cost, length and throughput of high-accuracy sequencing will also help. For example, sequencing larger portions of the rrn operon including some or all of the ITS and 23S genes may allow for serovar-level classification from single amplicon sequences (Graf 2021, Karst 2021), largely side-stepping the strain confusion problem. But key limitations will likely remain including a limit of detection based on relative abundance within the community of organisms being amplified.

The cost of the methodology we are proposing for the identification of serovars in isolates (barcoding) and in mixed community samples (metabarcoding) is critical to its practical importance. The price of traditional serotyping from isolates via molecular methods has been estimated to cost on average $42.37 per sample (Yoshida et al. 2016, Ford et al. 2021). Whole genome sequencing (WGS) based serotyping has been estimated to cost an average of $83.15 per sample, generally running at a higher cost than molecular serotyping (Ford et al. 2021). In the context of barcoding, i.e. molecular characterization from isolates, long-read amplicon sequencing costs of the full-length 16S rRNA gene are regularly advertised by Pacbio at $10 a sample, and more aggressive multiplexing with per-sample read targets of 200-300 reads/sample (sufficient to measure an isolate’s full 16S rRNA gene profile) could approach costs as low as $2/sample. However, those very low per-sample costs depend critically on high levels of multiplexing, and thus the need to regularly barcode hundreds or thousands of isolates. The financial aspects of applying our approach to metabarcoding samples is more difficult to ascertain, but it is fair to say that as the number of samples in which large amounts of non-target DNA might be present (such as food samples) the relative advantage of a targeted method like amplicon sequencing over untargeted sequencing increases.

Although real challenges exist to realizing the full potential of long-read amplicon sequencing of the 16S rRNA gene (and beyond), there is a clear value proposition going forward. Amplicon sequencing is an extremely effective approach for focusing sequencing on a taxonomic range of interest in the presence of large amounts of non-target DNA. New sequencing technologies are changing what is possible from standard methods like amplicon sequencing. This includes 16S rRNA gene sequencing, a specific (to prokaryotes) but broad (to bacteria) approach to characterizing complex microbial communities that is being supercharged by the high-accuracy long-read sequencing technologies that have been developed and are becoming increasingly available. As reference databases and sequencing technologies continue to progress, there are exciting new possibilities, including effective sub-species level classification from amplicon sequencing of the 16S rRNA gene across large swathes of pathogenic and non-pathogenic bacterial species.

## Acknowledgements

BJC, DOG, and ST were supported by USDA NIFA grant 2019-67021-29927. BJC was also supported by NIH grant R35GM133745. The funders had no role in study design, data collection and interpretation, or the decision to submit the work for publication.

## Author Contributions

DOG and BJC developed the method and algorithm for serovar prediction. DOG wrote the computational tool and code. LBH prepared isolate and environmental samples for analysis. BJC and ST conceptualized the research and acquired funding. DOG and BJC prepared the manuscript. All authors revised, edited, and assisted in writing the final version of the manuscript.

## Supplemental Tables and Figures

**Supplemental Table 1.** Serovar assignment outcomes for all *Salmonella* serovars containing at least 4 assemblies within our reference database. Correct, incorrect, and indeterminate rates are reported for each serovar. The test queries used to generate this data included all assemblies belonging to any serovar with at least 4 entries in our reference database.

| Serovar | Total in Group | Correct (%) | Indeterminate (%) | Incorrect (%) |
| --- | --- | --- | --- | --- |
| *Enteritidis* | 300 | 99.3 | 0 | 0.7 |
| *Typhimurium/Monophasic* | 288 | 95.1 | 2.4 | 2.4 |
| *Typhi* | 101 | 89.1 | 10.9 | 0 |
| *Infantis* | 60 | 98.3 | 1.7 | 0 |
| *Newport* | 54 | 88.9 | 1.9 | 9.3 |
| *Heidelberg* | 41 | 100 | 0 | 0 |
| *Kentucky* | 41 | 97.6 | 2.4 | 0 |
| *Anatum* | 36 | 94.4 | 5.6 | 0 |
| *Bareilly* | 34 | 85.3 | 8.8 | 5.9 |
| *Agona* | 33 | 97 | 3 | 0 |
| *Saintpaul* | 32 | 90.6 | 3.1 | 6.3 |
| *Indiana* | 29 | 79.3 | 10.3 | 10.3 |
| *Montevideo* | 29 | 96.6 | 3.4 | 0 |
| *Senftenberg* | 27 | 85.2 | 7.4 | 7.4 |
| *Hadar* | 26 | 100 | 0 | 0 |
| *Javiana* | 21 | 9.5 | 81 | 9.5 |
| *Muenchen* | 19 | 94.7 | 0 | 5.3 |
| *Dublin* | 18 | 83.3 | 0 | 16.7 |
| *Reading* | 18 | 72.2 | 22.2 | 5.6 |
| *Derby* | 14 | 57.1 | 21.4 | 21.4 |
| *Goldcoast* | 14 | 92.9 | 0 | 7.1 |
| *Rubislaw* | 14 | 71.4 | 28.6 | 0 |
| *Weltevreden* | 14 | 100 | 0 | 0 |
| *Schwarzengrund* | 12 | 100 | 0 | 0 |
| *Thompson* | 12 | 66.7 | 25 | 8.3 |
| *Albany* | 11 | 100 | 0 | 0 |
| *Choleraesuis* | 11 | 81.8 | 9.1 | 9.1 |
| *Bovismorbificans* | 10 | 80 | 0 | 20 |
| *Inverness* | 10 | 100 | 0 | 0 |
| *Tennessee* | 10 | 90 | 10 | 0 |
| *Gallinarum* | 9 | 55.6 | 0 | 44.4 |
| *Oranienburg* | 9 | 77.8 | 0 | 22.2 |
| *Gaminara* | 8 | 50 | 25 | 25 |
| *London* | 8 | 87.5 | 12.5 | 0 |
| *Braenderup* | 7 | 100 | 0 | 0 |
| *I 1,4,[5],12:b:-* | 7 | 14.3 | 28.6 | 57.1 |
| *Paratyphi A* | 7 | 100 | 0 | 0 |
| *Give* | 6 | 33.3 | 16.7 | 50 |
| *Mississippi* | 6 | 100 | 0 | 0 |
| *Brandenburg* | 5 | 40 | 20 | 40 |
| *Minnesota* | 5 | 40 | 20 | 40 |
| *Muenster* | 5 | 100 | 0 | 0 |
| *Uganda* | 5 | 100 | 0 | 0 |
| *V 66:z41:-* | 5 | 100 | 0 | 0 |
| *Worthington* | 5 | 100 | 0 | 0 |
| *Bredeney* | 4 | 0 | 50 | 50 |
| *II 1,4,12,[27]:b:[e,n,x]* | 4 | 50 | 50 | 0 |
| *IIIb 60:r:z* | 4 | 100 | 0 | 0 |
| *Mbandaka* | 4 | 0 | 0 | 100 |
| *Ohio* | 4 | 100 | 0 | 0 |
| *Panama* | 4 | 50 | 0 | 50 |
| *Paratyphi B* | 4 | 75 | 0 | 25 |

**Supplemental Figure 1.**
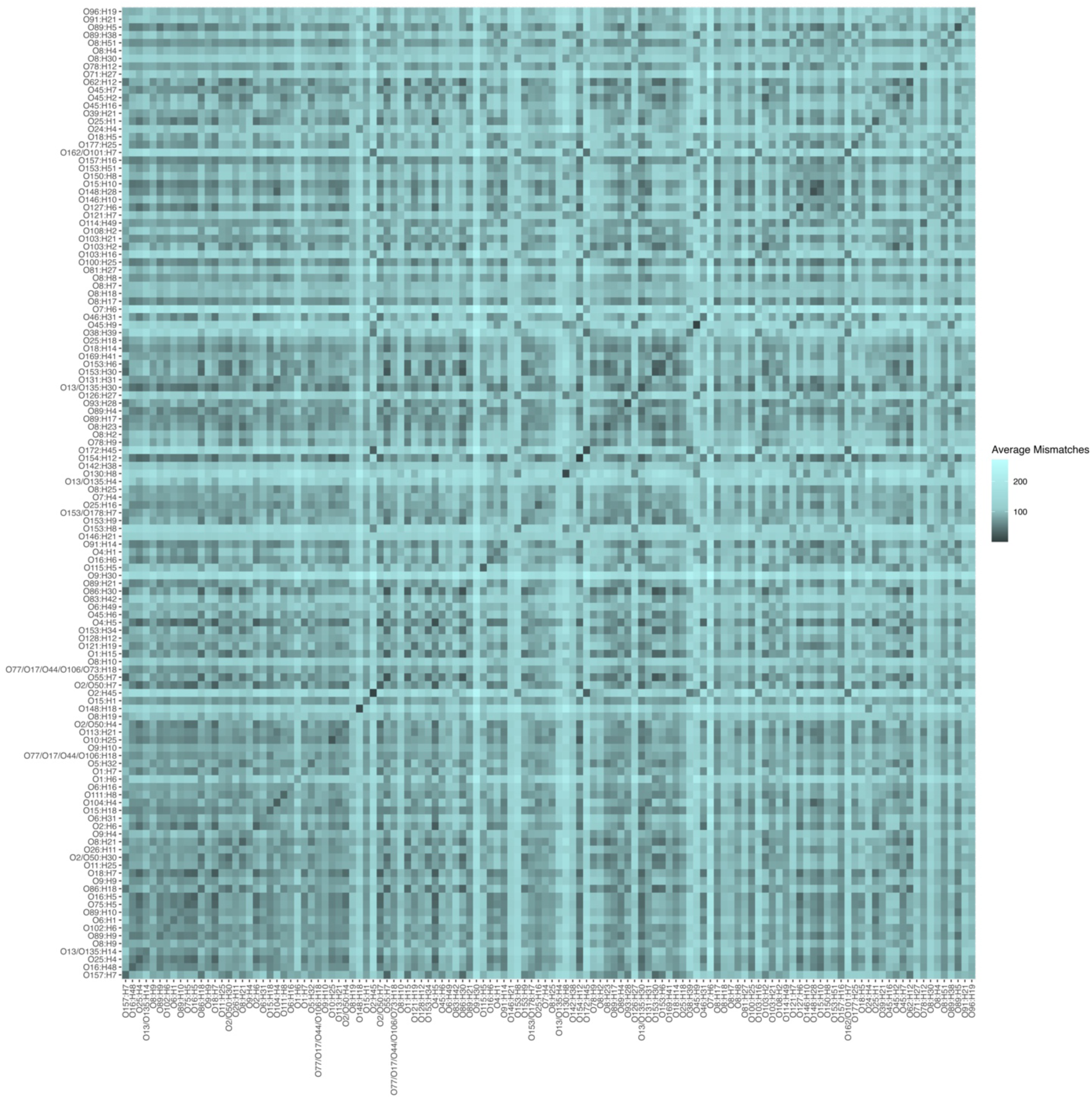
Average dissimilarity between the 16S rRNA gene profiles of *E. coli* serovars. Pairwise dissimilarities were defined as the number of nucleotide mismatches between alignments of the full set of 16S rRNA genes (the 16S rRNA gene profile) of *E. coli* assemblies, after optimally re-arranging the profiles relative to one another (Methods).

